# Frequency-dependent diffusion tensor distribution imaging in the evaluation of ischemic stroke

**DOI:** 10.1101/2025.05.19.654809

**Authors:** Sara Gröhn, Ángela Naranjo, Omar Narvaez, Maxime Yon, Buse Buz-Yalug, Santos Blanco, Daniel Topgaard, Esther Martinez-Lara, Ma Ángeles Peinado, Jussi Tohka, Alejandra Sierra

## Abstract

Non-invasive MRI is widely used to assess and monitor ischemic stroke, yet conventional approaches often lack sensitivity to subtle microstructural changes and struggle to evaluate tissue viability across lesion, penumbra, and distal regions. In this study, frequency dependent diffusion tensor distribution imaging (ωDTD) was combined with clustering of diffusion tensor distributions **D**(ω) and multivariate regression modeling to characterize ischemic tissue alterations in a whole brain section. Ex vivo ωDTD and histology were performed in rats subjected to middle cerebral artery occlusion (MCAO) or sham surgery (P = 17) 24 hours after reperfusion. Lesions showed cell loss and an increased presence of smaller, likely glial, cells. A random forest (RF) model was used to explain and predict histological parameters from diffusion tensor imaging (DTI), manually bin resolved ωDTD features, and cluster resolved ωDTD parameters. Model performance was evaluated using leave one animal out cross validation (LOO CV). ωDTD features better represented cell number than DTI metrics (ωDTD R^2^ = 0.73 vs. DTI R^2^ = 0.49), with similar advantages for nuclear area and circularity (ωDTD R^2^ = 0.64 and 0.61 vs. DTI R^2^ = 0.40 and 0.35). The RF model further proved beneficial in capturing complex, nonlinear relationships between MRI parameters and tissue characteristics. Overall, these results indicate that ωDTD provides richer microstructural information than standard DTI, and that combining ωDTD with advanced machine learning methods enhances interpretation of ischemic tissue damage.

## INTRODUCTION

Ischemic stroke is an significant health and economic burden on a global scale, with up to 7.6 million incident cases annually [1]. During the ischemic event, the brain undergoes a series of complex biochemical and structural changes such as shifts in tissue composition and neuronal damage. Ischemic stroke arises from obstructed blood flow in a section of the brain leading to a lack of oxygen and nutrients in the affected area. This results in cellular injury and eventually cell death if blood flow is not restored [2, 3]. While traditional magnetic resonance imaging (MRI) techniques used in clinic e.g. T2-weighted imaging, diffusion MRI (dMRI), diffusion tensor imaging (DTI) and perfusion weighted imaging (PWI), are effective in detecting large-scale structural damage in the ischemic lesion core, the viability of the tissue in the core, penumbra, and distal areas to the lesion often remains undefined based on these methods [4, 5, 6, 7, 8]. These methods extract rather limited tissue information during pathological processes and often fail to detect subtle changes crucial to understand tissue viability or treatment responses after reperfusion. Therefore, Ischemic stroke requires novel solutions to these limitations.

Diffusion tensor distribution imaging (DTD) is an emerging MRI method with the potential to provide a solution to the challenges posed by conventional MRI. The advanced acquisition protocols of DTD utilize tensor valued diffusion encoding. This allows sub-voxel parameter estimation of cellular features such as “size”, “shape” and orientation, enabling a more nuanced and detailed analysis of tissue microstructure [9, 10]. Additionally, frequency-dependent DTD (*ω*DTD) exploits frequency dependence of diffusion encoding to estimate the effect of restriction [11, 12]. Restriction describes how the diffusion of water is constrained by the barriers of different tissue microenvironments, such as cellular membranes, at a few micrometers scale depending on the diffusion encoding frequency range. It can therefore provide more detailed insights into tissue composition and integrity for the targeted feature scale [13, 10, 14]. The parameters diffusivity, micro-anisotropy, and orientation more familiar from the literature [15, 9], are calculated from the frequency-dependent diffusion tensor distribution **D**(*ω*) within each voxel. These parameter distributions can then be manually segmented into three bins to discriminate the specific signal-fractions within the voxel, such as those specific of white matter (WM), grey matter (GM) and free water (FW) like tissues [16, 10]. This simple binning operation based on two dimensions only allows computing specific parameter maps taking advantage of the correlation between all parameters [12, 9]. However, even in a healthy brain these manually divided “bins” can not accurately represent the complexity of the tissue and especially in pathological conditions, the binning becomes all the more challenging to interpret. To utilize more than two dimensions in the binning operation, we recently introduced Gaussian mixture-model based clustering of the tensor-valued distributions [17]. This approach allows the unsupervised and automated division of the per-voxel parameter distributions into k cluster-resolved fractions instead of three manually divided bin-resolved fractions. The additional solution space subsets produced by the clustering could potentially provide increased specificity in defining the ischemia related changes with MRI.

In this study, we combined *ω*DTD, unsupervised clustering, and a multivariate statistical model to understand the relationship between MRI and histology following ischemic stroke. Our approach utilises the power of non-linear regression, specifically random forests (RF), to explain and predict ischemic damage related tissue changes at 24-hours after reperfusion. Additionally, we investigate how the prediction accuracy depends on the input MRI parameters. We evaluated the microstructural changes related to ischemia based on altered cell morphology in the whole brain slice and included four different MRI parameter sets ranging from conventional DTI parameters to *ω*DTD cluster-fraction parameters. This study brings novelty to the field in specifically concentrating on exploring the relation between *ω*DTD and histological changes in ischemic stroke.

## MATERIALS AND METHODS

### Animals and stroke model

Seventeen adult male Wistar rats (250-450 g; Charles River, Wilmington, MA, USA) were used in this study. The rats were housed individually in a climate-controlled room under 12/12-hour light/dark cycle on an ad libitum diet. A transient ischemic stroke was induced in the right hemisphere using the middle cerebral artery occlusion (MCAO) model [18]. The MCA was occluded with an intraluminal filament (Doccol Corporation, Redlands, CA, USA) inserted through the right common carotid artery for 45 minutes before reperfusion as described in [19, 20]. The animals were divided into three groups. The first MCAO group received empty nanoparticles (NP) (MCAO+∅NP, n = 6). The second MCAO group received NPs filled with neuroglobin (MCAO+Ngb-NP, n = 6), an endogenous oxygen-binding globin protein found to play a neuroprotective role against many brain injuries, including ischemia [18, 21, 22]. The NPs were injected through the tail vein. The third group was sham-operated, and also received empty NPs (sham, n = 5) to maintain experimental consistency and eliminate confounding variables.

At 24-h post-reperfusion, the animals were deeply anesthetized for a transcardial perfusion (left ventricle) using 5% isoflurane in O_2_/N_2_ mixture. This time point was selected to capture acute stroke pathology. Previous studies have shown that MCA occlusion in rodents produces peak infarct volume (assessed by TTC staining) at 24 h post-reperfusion, which remains stable through day 7, while apoptosis peaks at 24–48 h [23, 24, 25]. The perfusion was performed first with 150 ml 0.01 M, pH 7.4 phosphate-buffered saline (PBS) for 5 minutes at 30 ml/min and continued with 200 ml 4% paraformaldehyde (PFA) and 10 u/ml of heparin for 7 minutes also at 30 ml/min. Then, finished with 300 ml 4% PFA for 20 minutes at 15 ml/min. After perfusion, the brains were carefully extracted from the skull and post-fixed in the same solution for 4 h, before being placed 4% PFA until imaging. The animal model was provided and supervised by the Animal Production and Experimental Centre and the Ethical Committee for Animal Experimentation (CEEA, approvals #23/05/2016/090 and #10/02/2021/011) at the University of Jaén, Spain. All procedures were performed in accordance with Spanish legislation (law 32/2007, amended by law 6/2012 and by the Royal Decree 53/2013) and European Union Directive guidelines 2010/63/EU.

### *Ex vivo ω*DTD acquisition and data processing

*Ex vivo* MRI data was acquired in a vertical 11.7-T Bruker Avance-III HD spectrometer equipped with a MIC-5 probe (on-axis 3 T/m maximum gradient amplitude) and 10-mm volumetric coil. The brains were divided into two hemispheres and the right hemisphere containing the ischemic lesion was imaged. The hemispheres were then immersed in gadoteric acid (0.0005 mmol/ml, Clariscan, GE HeatlhCare) in PBS pH 7.4 for 24 h prior to imaging to recover longitudinal and transverse relaxations and improve SNR for ex vivo [26]. Before MRI, the hemispheres were placed inside a 10 mm polyethylene tube and immersed in a perfluorinated compound (Galden, Solvay, Italy) to effectively suppress the background signal.

MRI data was obtained with a multi-slice multiecho (MSME) 2D sequence customized for tensor-valued diffusion encoding. The sequence included variable b-values (700 - 8000 s/mm2), normalized anisotropy bΔ (−0.5, 0, 0.5 and 1), TR = 800 ms, TE = 30 ms, orientation (*θ,ϕ*) and centroid frequency *ω*_cent_/2*π* at 10% and 90% percentile (34 - 115 Hz). The images were acquired at approximately −0.60 mm from bregma with an in-plane voxel resolution of 80 x 80 *µ*m2, slice thickness of 250 *µ*m and a matrix size of 150 x 150 x 1. In addition, anatomical data of the same anterior-posterior location was acquired using the rapid acquisition with relaxation enhancement with variable repetition time TR (RARE VTR) sequence to reduce scan time and sample the longitudinal relaxation more effectively. The total acquisition time was 4 hours and 12 minutes. After scanning, the brains were stored in 2% PFA solution to await histology processing.

The images were reconstructed using Bruker’s ParaVision 6.0.1 [27] into real and imaginary domains and separately preprocessed using 1) a python-based denoising algorithm [28] based on the exploitation of data redundancy in principal component analysis domain, and 2) a MRtrix3 based Gibbs-ringing removal algorithm [29] based on local sub-voxel shifts. After preprocessing the real and imaginary domains were used to obtain the magnitude image.

### Estimation of nonparametric D(*ω*) distributions

The nonparametric frequency-dependant diffusion distributions **D**(*ω*) were derived through approximating the **b** encoded signal S as shown in [30]

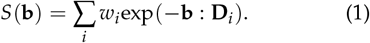

The *ω*-dependent self diffusion tensor **D**(*ω*) is acquired by inverting equation (1) and approximating the **D**_*i*_(*ω*) as an axisymmetric tensor,

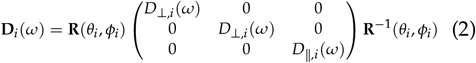

where **R**(*θ*_*i*_, *ϕ*_*i*_) is a rotation matrix and parallel and perpendicular eigenvalues *D*∥ _,*i*_(*ω*) and *D* ⊥_,*i*_(*ω*) are given by Lorentzian transformations [30] at the frequencies Γ*∥*/*⊥,i* according to

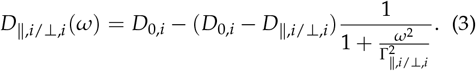

Using the axisymmetric Lorentzian approximation, the components of the **D**(*ω*) distribution are described by the weight *w* and parameter set [*D*_A_, *D*_R_, *θ, ϕ, D*_0_, Γ_A_, Γ_R_,]. The data inversion is then performed using an extension of the Monte Carlo algorithm with bootstrapping as described in [12, 30, 31] using the md-dmri MATLAB toolbox [32].

The bootstrapping consisted of 100 repetitions using random sampling with replacement for 8 output components. The data inversion yielded an ensemble of 100 independent solutions of the **D**(*ω*) per-voxel, each consisting of the 8 output components; weights and coordinates (*w*, [*D*_A_, *D*_R_, *θ, ϕ, D*_0_, Γ_A_, Γ_R_,]), within the pseudo-randomly sampled analysis space. The inversion limits were *D*_R/A/0_ ∈ [0.005, 5] · 10−9 and Γ_R/A/0_ ∈ [0.000001, 1] ·105. Then the more familiar and meaningful parameters of isotropic diffusivity *D*_iso_, normalized anisotropy *D*_Δ_ and normalized squared anisotropy 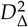 were calculated through

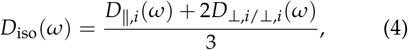

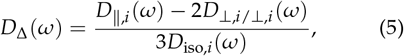

and

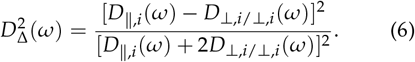

The effect of restriction on *ω*-dependent *D*_iso_ and 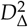 has been explored in oscillating gradient encoding [33, 34] and other *ω*DTD [12, 35, 30, 36, 37] studies and can be quantified as

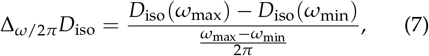

and

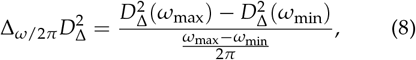

where *ω* is valued from *ω*_min_ to *ω*_max_ within the investigated frequency window (10% to 90%). The distributions can then be visualized as contour plots in a 2D 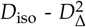 space as well as bin-resolved signal fractions *f*_bin*n*_, means E_bin*n*_ [·], variances V_bin*n*_ [·] and covariances C_bin*n*_ [·, ·] from which the per-voxel statistical descriptors values were obtained to display the quantitative parameter maps [9]. Here we have only considered the bin-resolved signal fractions *f*_bin*n*_, means E_bin*n*_ [·] calculated as

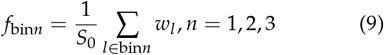

and

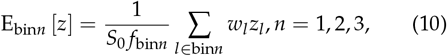

where *z* represents *D*_iso_, 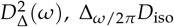 and 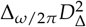. Subscript *l* refers to the bootstrap samples and *S*_0_ is the non-diffusion weighted image.

Additionally, basic DTI maps of mean diffusivity (MD) and fractional anisotropy (FA). These were derived from the diffusion tensor [38], calculated based on the orientation information in the *ω*DTD data.

### Clustering of D(*ω*) distributions

The traditional manual binning is performed based on predefined limits of the isotropic *D*_iso_ and anisotropy 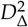 space as follows; bin 1: *D*_iso_ ∈ [0,1] and 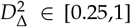, bin 2: *D*_iso_ *∈* [0,1] and 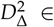 [0,0.25] and bin 3: *D*_iso_ ∈ [1,2] and 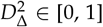. The bin-resolved maps are produced based on this division. In turn, the clustering of the bootstrapped *D*_iso_, *D*_Δ_ - for accessing information on prolate and oblate tensor shapes [39, 40] - and Δ_*ω*2*π*_ *D*_iso_ per-voxel distributions was performed as described in [17] using Gaussian mixture model clustering. The clustering was performed with *k* = 2, …, 8 to avoid predefining the number of clusters. BIC (Bayesian information criteria) was used to aid in the selection of optimal k (see supplementary figure S8). The spatial information from the clusters *clus*_n_ can be visualised with the per-voxel normalized frequency maps *F*_*clus*n_, *n* = 1, 2, 3, …, *k*, where *k* is the number of clusters. Then, for the optimal *k* value, we produced the distributions and cluster-resolved signal fraction *f*_clus_ maps [17]. The cluster-resolved signal fractions can be obtained following a similar formulation as with the bin-resolved signal fractions. These cluster-resolved signal fractions *f*_clus_ can then be visualized using contours in the 2D 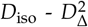 space and per-voxel means E_clus*n*_ [·] parametric maps calculated as

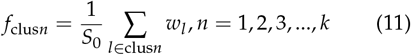

and

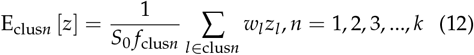

### Histology

The hemispheres were first immersed in a cryoprotectant solution of glycerol 20% in 0.02 M potassium PBS (KPBS) pH 7.4 for 48 h before being frozen in dry-ice and stored at −80°C. Before sectioning, the brains were embedded in optimal temperature cutting medium. The sectioning was performed in the coronal plane at 30-*µ*m slice thickness using a cryostat (Leica CM1950, Leica Microsistemas S.L.U., Spain). The sections were then stored in a cryoprotectant tissue-collecting solution (30% ethylene glycol, 25% glycerol in 0.05 M sodium phosphate buffer) at −20°C until staining. For the Nissl staining, the sections were washed with 0.1 M PB (pH 7.4), then incubated in a chloroform and absolute ethanol (1:1) mixture. The sections were hydrated in descending concentrations of alcohol: absolute ethanol twice for 2 minutes; 96% for 2 minutes; 70% for 2 minutes; and 50% for 2 minutes. Then, the sections were distilled in water for 1 minute. Then, the sections were immersed in 0.125% thionine to assess both tissue damage severity and cytoarchitectonics until the appropriate coloring was achieved. The excess thionine was washed off with two washes using distilled water and then the sections were dehydrated increasing alcohol concentrations: 50% for 2 minutes; 70% for 2 minutes; 96% for 2 minutes and absolute ethanol twice for 2 minutes. Finally, 3 x 5 minutes washes were performed using xylol before mounting the sections in dibutylphthalate polystryrene xylene. The Nissl-stained sections were scanned at 20x magnification using a BX dotslide digital virtual microscope equipment, Olympus, at the Scientific Instrumentation Center of the University of Granada, Spain.

### Non-linear regression and analysis

To assess the relationship between MRI and histology, we performed a non-linear regression analysis using the RF algorithm [41]. A total of *P* = 11 animals were included. From the original sample size of 17, 1 brain per group was allocated for serial-block electron microscopy (SBEM) analysis and therefore had an MRI acquisition not comparable with the MRI sequence described in this paper. Additionally, histology was performed for only one sham animal. For the regression analysis, we extracted quantitative parameter maps from the histology and selected the photomi-crographs (0.3231 x 0.3231 *µ*m2) that gave the closest match to the MRI slices (approx −0.60 mm from bregma) for analysis. The photomicrographs were then divided to 80 x 80 *µ*m2 tiles before applying the QuPath watershed segmentation technique-based cell detection [42]. From the resulting cell detection measurements, we reconstructed three quantitative parameter maps depicting the number of cells per tile as well as the average nucleus area and circularity per tile. These histological parameter maps were then masked and downsampled using nearest neighbour interpolation. Lastly, the histological parameter maps were registered with the DTI FA map of the corresponding brain using a rigid similarity transform optimized by MATLAB’s imregtform function. The registration minimized the negative mutual information [43] via the One-Plus-One Evolutionary (OPOE) algorithm [44, 45] suited for multimodal images. The maps were then prepared for the regression analysis by flagging possible negative (errors) and zero-valued voxels within the brain tissue as missing values. The pipeline is outlined in **Figure 1**.

**Figure 1:**
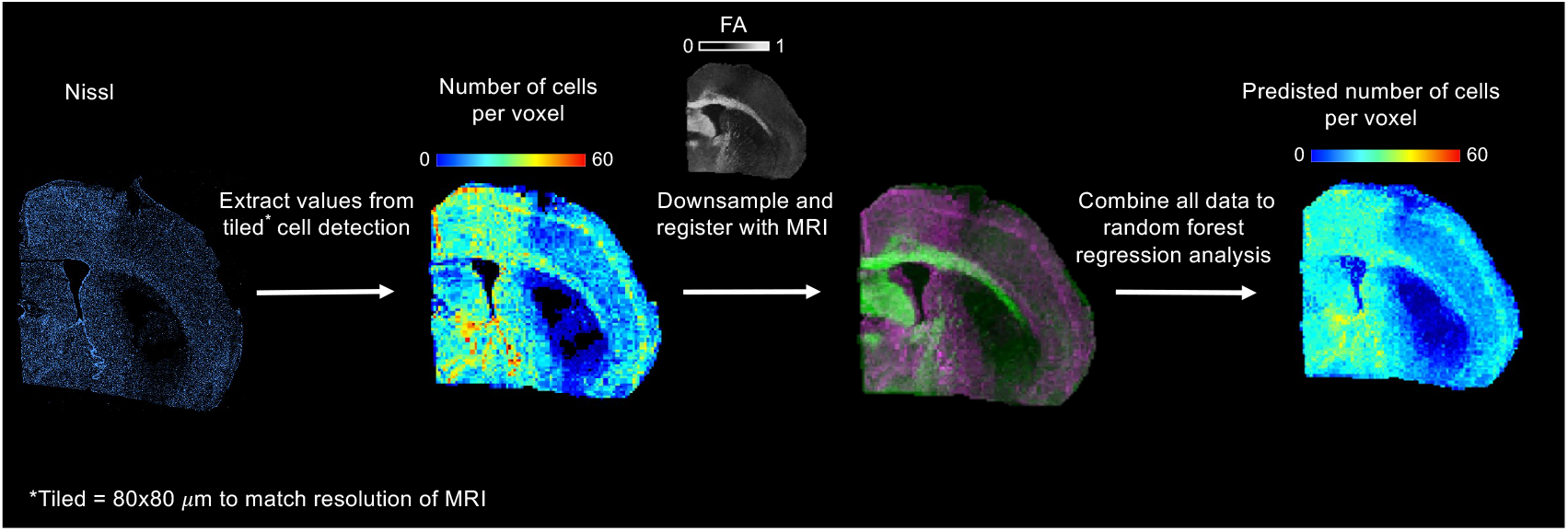
Overview of the non-linear regression analysis pipeline. For each brain, the Nissl-stained photomicrographs were tiled to 80 x 80 µm2 sections to match the resolution of the MRI data. Cell detection was performed on these images to extract quantitative information. Tiled histological parameter maps were then reconstructed, downsampled and individually registered with the corresponding DTI FA map. For the non-linear regression analysis, MRI parameters were grouped as follows: (1) conventional MRI consisting of T2-weighted, MD and FA maps; (2) ωDTD per-voxel mean and rates of change per frequency maps; (3) ωDTD bin-resolved mean and rates of change per frequency maps; and (4) ωDTD cluster-resolved mean and rates of change per frequency maps. For each MRI parameter set, all brains were combined into a single dataset, which was used to predict the histological parameters using a with 1000 trees.

The MRI data was then divided into four sets to evaluate, which one best predicted the histological parameters. The sets were: (1) conventional DTI-MRI consisting of MD and FA maps (2 maps); (2) *ω*DTD per-voxel mean and rates of change per frequency maps (4 maps: per-voxel mean D_iso_, 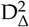, Δ_*ω*2*π*_E[*D*_iso_] and, 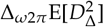; (3) *ω*DTD bin-resolved mean and rates of change per frequency maps (12 maps: bin-resolved D_iso_, 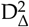, Δ_*ω*2*π*_E[*D*_iso_] and, 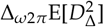); and (4) *ω* DTD cluster-resolved mean and rates of change per frequency maps (28 maps: cluster-resolved D_iso_, 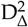, Δ_*ω*2*π*_E[*D*_iso_] and, 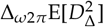). The MRI data was prepared for the regression analysis as follows: (1) Individual maps were masked and processed for outliers by using the third quartile to remove such as, the bright halo effect around DTI MD maps. (2) The MRI datasets were scaled using z-score normalization. This preprocessing was done to help optimize the data for the RF regression task [46].

The RF analysis was performed using the Tree-Bagger function by MATLAB with 1000 trees. The predictor data matrix was built as

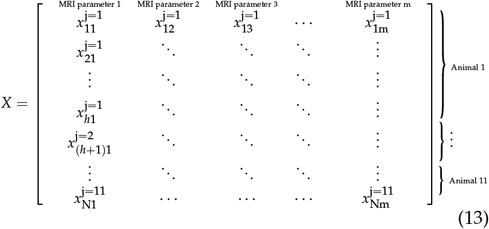

where *m* is the number of MRI parameters per set and the MRI maps from all animals *j* were concatenated row-wise, with the number of datapoints all together being N. The target *Y* (Nx1) was built following the same logic as *X*

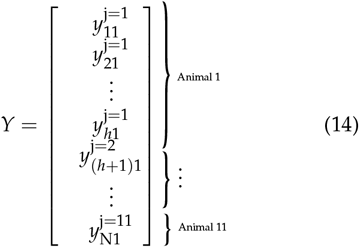

A separate model was trained for each individual histological and MRI set combination. Based on the trained RF models, we predicted the histological parameter maps.

The performance of the RF models was assessed using leave-one-animal-out cross-validation (LOO CV) as in [47] using the mean (E[·]) and variance (V[·]) of the different metrics:

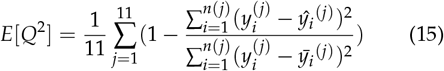

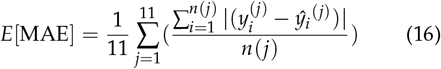

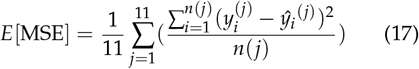

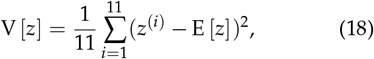

where 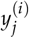 is the true histological parameter of animal *j* at the voxel *i, n*(*j*) is the number of data-points for animal *j*, 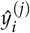 is the predicted histological parameter and 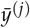 is the average of the true histological values. The *Q*2 metric sheds light on the model’s ability to generalize as it provides an internal measure for consistency between CV predictions and original data [48], while MSE and MAE tell of the accuracy and generalizability of the model as well as potential overfitting [46, 49]. The *R*2 [50] was used as a performance measure to estimate the goodness-of-fit for the model. The CV analysis was performed using an in-house MATLAB code, available at https://github.com/jussitohka/mrihistology.

To compare the predictive performance of the different MRI feature sets, we used a paired permutation test (100,000 permutations) on the subject-wise performance metrics (*R, Q*2, MSE, and MAE). For each histological parameter and performance metric, we treated every MRI method–histology combination as a separate model and then tested all pairwise differences between these models (e.g. DTI–cellularity vs *ω*DTD per-voxel mean and rates of change with frequency maps–cellularity). For a given metric and histological parameter, the test statistic was the mean within-subject difference between two methods (e.g. *R* for DTI minus *R* for *ω*DTD per-voxel mean and rates of change with frequency maps). This approach avoids distributional assumptions such as normality of differences and is well suited to the relatively small sample sizes and potentially non-Gaussian performance metrics common in imaging-based prediction studies.

Shapley additive explanations (SHAP) were calculated with the shapley and fit functions in MATLAB and used to obtain local feature attributions for a subset of 40 voxels per animal across cross-validation folds. This subset size was chosen to balance representativeness with the substantial computational cost of SHAP estimation, and voxels were strategically selected to capture both lesioned and adjacent tissue. Local Shapley values were then aggregated across query points, folds, and animals to derive global feature importance, following the framework proposed for tree ensembles by [51], where many local explanations are combined to characterize global model behavior.

## RESULTS

### Qualitative ischemic changes in conventional- and bin-resolved *ω*DTD parameter maps

**Figure 2A** and **D** show a conventional T2-weighted anatomic map along with DTI parameter maps mean diffusivity (MD) and fractional anisotropy (FA) for representative sham-operated and MCAO rats. The anatomic T2-weighted map of the MCAO brain (**Figure 2D**) shows a slightly increased signal intensity within the lesion, which is consistent with increased presence of water in the region. Both the MD and FA maps, in turn, show a decrease in signal values in the lesion. The MD map provides a visually strong contrast for the lesion with clearly delineable borders.

**Figure 2:**
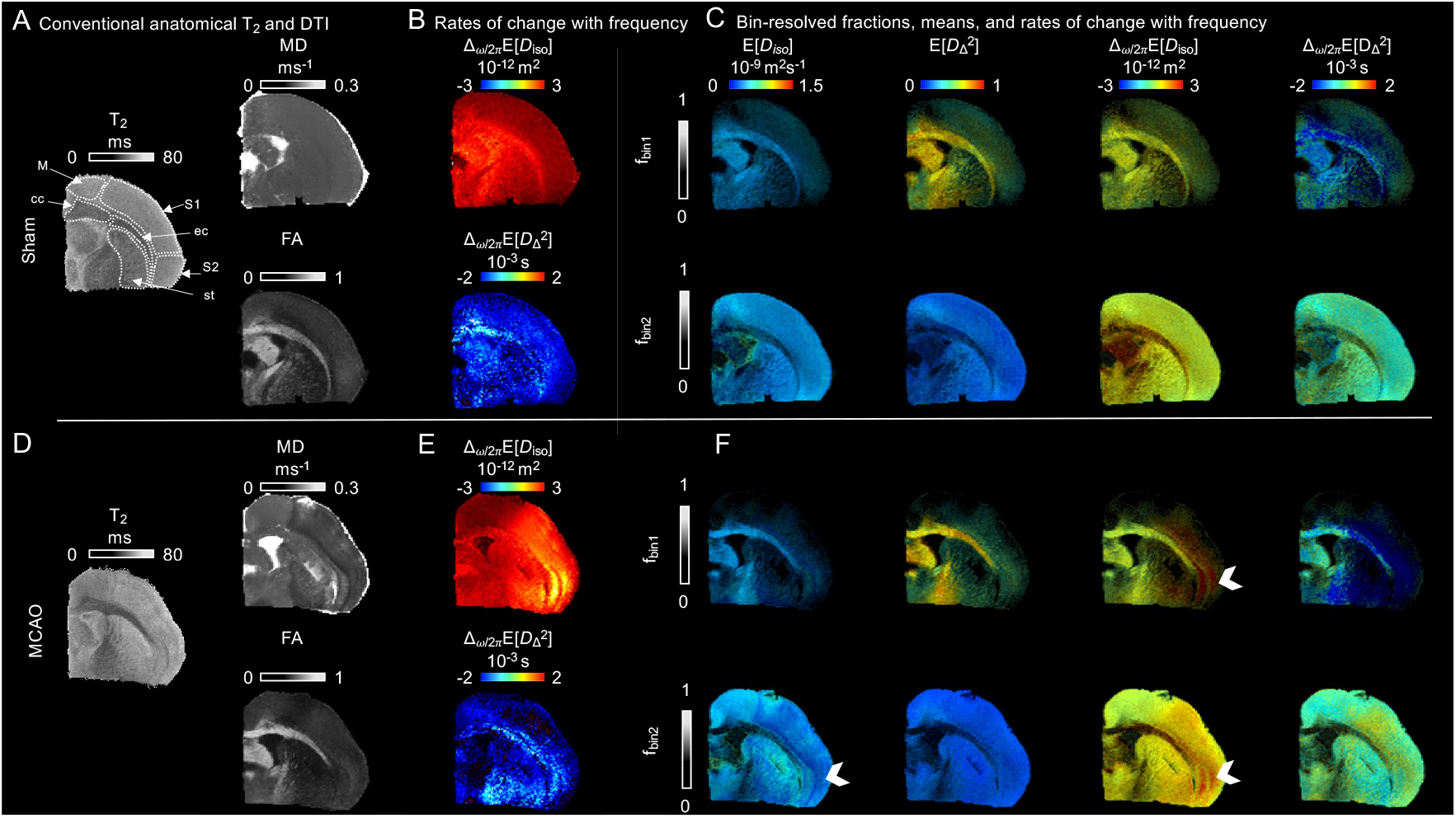
Conventional anatomical T2 and DTI metrics map compared to ωDTD parameter maps for representative sham and MCAO brains. **(A)** and **(D)** Conventional T2-weighted anatomic maps together with classic DTI parameter maps of MD and FA. **(B)** and **(E)** ωDTD maps showing per-voxel rates of change with frequency, along with **(C)** and **(F)** bin-resolved parametric maps for manually defined bin limits in the same representative brains (**equations** (4),(6), (7), **and** (8)). The brightness of the bin-resolved maps reflects signal fraction, while the color indicates parameter values. The ωDTD maps were computed using the ωDTD distribution (**equation** (2)) at ω/2π = 34.4 Hz and the rates of change with frequency maps were calculated with ω/2π = 34.4-115 Hz. The labels are M = motor cortex, S1 = primary somatosensory cortex, S2 = Secondary somatosensory cortex, cc = corpus callosum, ec = external capsule, and st = striatum. The arrows indicate the lesion in the somatosensory cortex.

The *ω*DTD parameters display multiple contrasts and highlight different regions of the lesion depending on the parameter map. **Figure 2B** and **E** show the rates of change with frequency maps Δ_*ω*2*π*_E[*D*_iso_] and 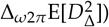 derived from the **D**(*ω*) distributions. The classic figure for displaying the *ω*DTD parameter maps, as in e.g. [12, 30], can be seen in supplementary figure S9) for a representative sham brain. The Δ_*ω*2*π*_E[*D*_iso_] map shows excellent lesion contrast across all the MCAO brains. The bin-resolved fractions, means and rates of change per frequency maps in **Figure 2C** and **F** depict the traditional manual division of the **D**(*ω*) distributions into “bins”. We present these classical bin-resolved maps as a qualitative point of reference for the clustering-based approach.

The MCAO frequency-dependent Δ_*ω*2*π*_E[*D*_iso_] map shows visually a increased signal in the somatosensory cortex as well as the striatum. The 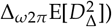 map also shows an increase in values in those same areas. The 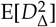 map of *f*_bin1_ shows increased values in the external capsule indicating more uniform orientation. In turn, the E[*D*_iso_] of *f*_bin1_, shows signal decrease in the external capsule denoting slower diffusion. Both maps also demonstrate changes in the somatosensory cortex when compared qualitatively to the representative sham-operated rat in **Figure 2B**. The Δ_*ω*2*π*_E[*D*_iso_] map of *f*_bin1_ shows an apparent increase in the values in the external capsule, striatum and somatosensory cortex (blue arrows). The E[*D*_iso_] map of *f*_bin2_ delineates visually very clear lesion borders similar to the MD map of **Figure 2D** and along with the 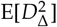 map of *f*_bin2_ shows hypointensity in the somatosensory cortex as well as the striatum. The Δ_*ω*2*π*_E[*D*_iso_] and 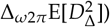 maps of *f*_bin2_ show increased signal values in those same regions.

### Clustering of D(*ω*) distributions

The parametric distributions of *D*_iso_, and 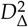 are presented in Fig. 3 for representative sham-operated and MCAO brains derived from the unsupervised clustering of the (D)_*ω*_, distributions. The optimum *k* based on the BIC (see supplementary figure S8) and [17] was 7. Using *k* = 7, the clustering consistently produced distributions presenting identifiable characteristics for each cluster-fraction (*f*_clust_) as can be seen in **Figure 3**. The first cluster-fraction is a WM-like fraction (*f*_clust1_), with *D*_iso_, and *D*_Δ_ distributions depicting low diffusivities, along with high anisotropy, diffusion characteristics typical in WM-like tissue, such as the corpus callosum and external capsule, but also including WM in cortical layers. The second cluster-faction is a *D*_Δ_ = −0.5 fraction (*f*_clust2_), followed by fraction of oblate shapes sensitive to frequency (*f*_clust3_). These first three E[*D*_iso_] cluster-fraction distributions for the MCAO also displayed a two peaked distribution in comparison to the single peaked appearance of the distributions in the sham-operated brains (black arrows in **Figure 3**).

**Figure 3:**
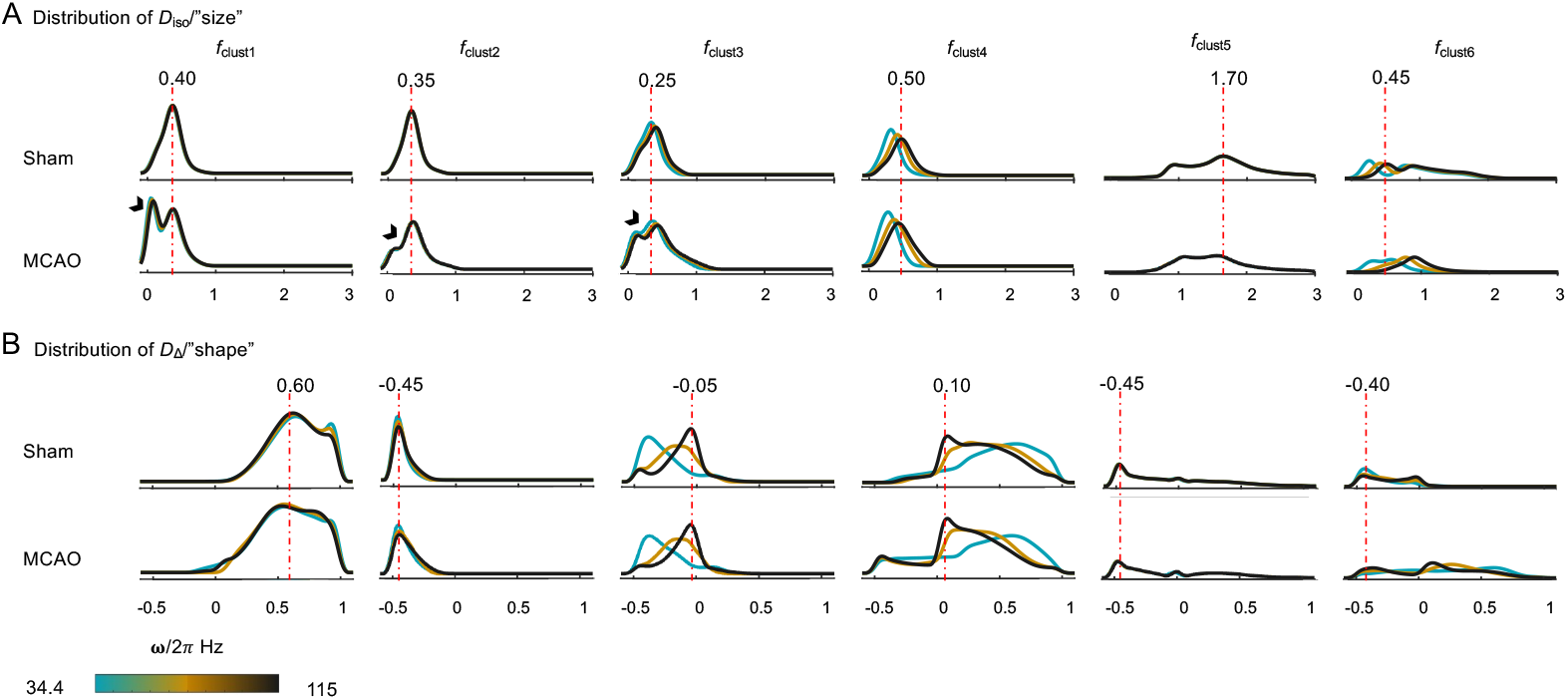
Distributions of D_iso_ and D_Δ_ for the cluster-fractions for representative sham and MCAO brains. The dotted red lines indicate the more prevalent value of D_iso_ and D_Δ_ for the sham-operated brain. Different frequencies are represented by colors 34.4 ω/2π Hz in blue, 74.7 ω/2π Hz in brown and 115 ω/2π Hz in black. The arrows point to the dual peak appearance of the D_iso_ distribution of the MCAO brain.

The fourth and sixth cluster-fractions (*f*_clust4_ and (*f*_clust6_) were characterised by high diffusivities and a widespread in the *D*_Δ_ distributions along with a dependency on frequency, indicating restriction. The fifth cluster-fraction is a FW-like fraction (*f*_clust5_) defined by with high diffusivity and low anisotropy values. And finally, a last spurious cluster-fraction (*f*_clust7_) also appeared consistently for all brains and is omitted from all analyses. For plotting the frequency dependency of these parameters, we used centroid frequencies 34.4, 74.7 and 115 *ω*/2*π* Hz.

Figure 4. shows the cluster-resolved mean parametric maps for representative sham-operated and MCAO brains derived from the unsupervised clustering of the (D)_*ω*_, distributions using *k* = 7. All the cluster-fractions, apart from the FW-like fraction (*f*_clust5_), showed distinct lesion contrast. While the cluster-resolved maps of E[*D*_iso_] showed a decrease in values in the lesion for the first three fractions, the lesion borders vary between the maps. The *f*_clust1_ depicts the changes in WM-like tissues, and while the different parameter maps show minor changes in the external capsule, the corpus callosum remains unchanged when compared to the corresponding sham-operated maps (**Figure 4A**). These *f*_clust1_ maps also show changes in the WM-like structures in the cortex, e.g. cortical axons. The 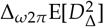 map of *f*_clust1_ shows prominent contrast for the lesion in of the somatosensory cortex.

The E[*D*_iso_] map of *f*_clust2_ depicts a diffuse and irregular lesion boundary, especially between the motor and somatosensory cortex. In comparison, the lesion boundaries in the E[*D*_iso_] map of *f*_clust1_ and *f*_clust3_ are distinct. The *f*_clust4_ and *f*_clust6_ demonstrate sensitivity to the lesion specifically, with *f*_clust6_ showing primarily the lesion and the visibility of the surrounding brain structures is minimal (white arrows in **Figure 4B**). The E[*D*_iso_] of *f*_clust4_ in turn shows hyperintensity in both primary and secondary somatosensory cortex, while *f*_clust6_ predominantly has signal only the secondary somatosensory cortex, the epicentre for the most intense damage. However, when comparing the lesion boundaries within the *f*_clust4_ and *f*_clust6_, there is a correspondence across the parameters. In turn, the 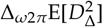 of *f*_clust4_ displays decreased signal values inside the lesion region. Additionally, Δ_*ω*2*π*_E[*D*_iso_] shows high values in *f*_clust4_ and *f*_clust6_ indicating high values of restriction.

**Figure 4:**
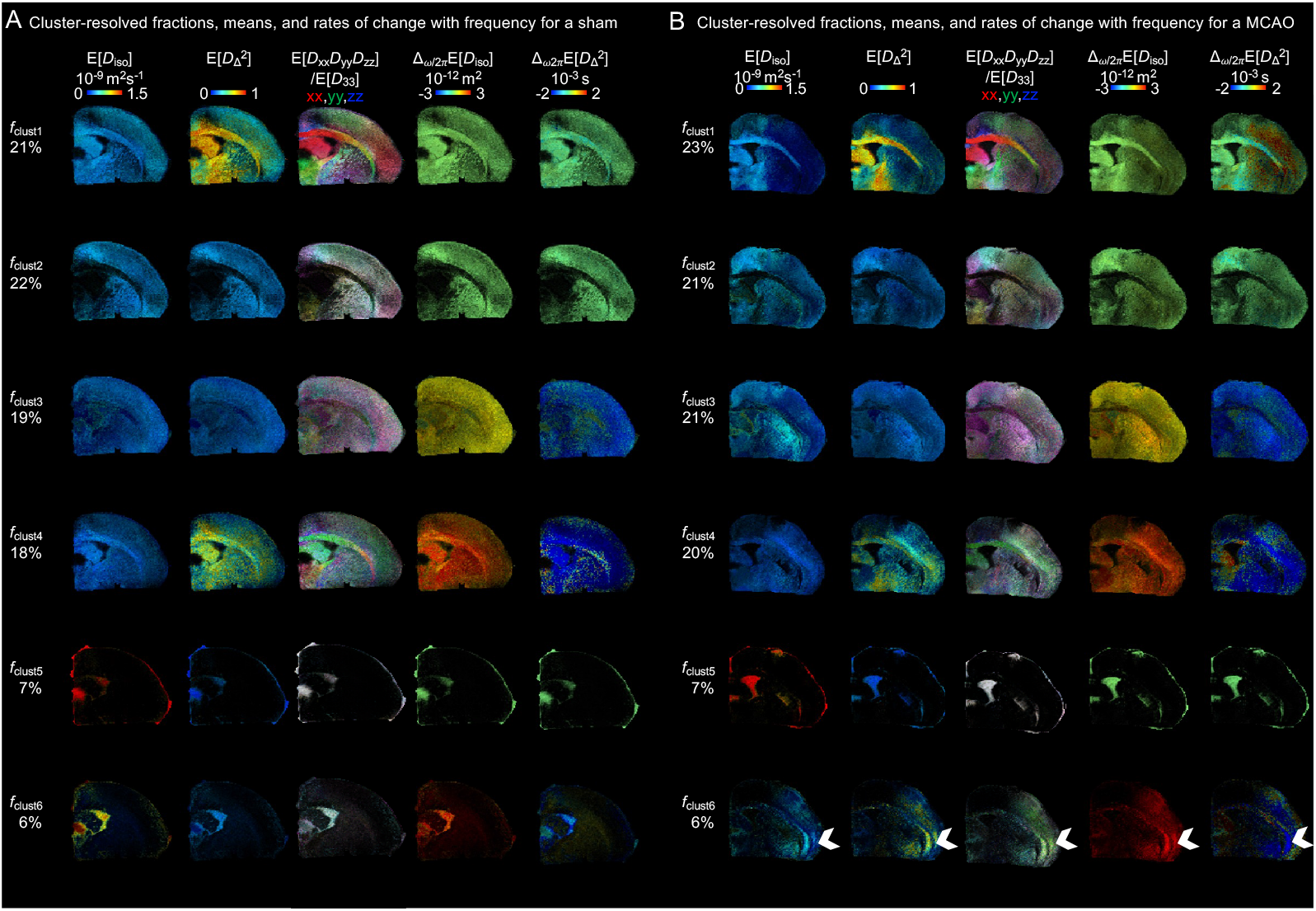
Cluster-resolved ωDTD parameter maps derived from D(ω) distributions for representative sham and MCAO brains. **(A)** and **(B)** Cluster-resolved ωDTD maps showing the cluster-fraction, with percentages indicating the per cluster normalized frequency in representative sham and MCAO brains. The color of the cluster-resolved maps indicates parameter values. The ωDTD maps were computed using the **D**(ω) distribution (eq. 2) at ω/2π = 34.4 Hz and the rates of change with frequency maps were calculated with ω/2π = 34.4-115 Hz. The arrows indicate the lesion in the somatosensory cortex.

### Relationship between histological and MRI parameters

A photomicrograph of a Nissl-stained section from a representative MCAO brain is shown in **Figure 5A**. The lesion was primarily located in the somatosensory cortex and was characterized by extensive cell loss and a presence of smaller cells that are most likely glial cells. The boundary of the lesion in the motor and primary somatosensory cortex (S1) demonstrated a clear transition between adjacent and damaged tissue, highlighting the regionality of the ischemic damage. The lesion boundary in the primary somatosensory cortex is reflected in the MRI parameter maps. The striatum (st) exhibited a significant loss of structures. The corpus callosum (cc) and the motor cortex (M) (see **Figure 5A**) were intact, exhibiting normal cellular density.

**Figure 5:**
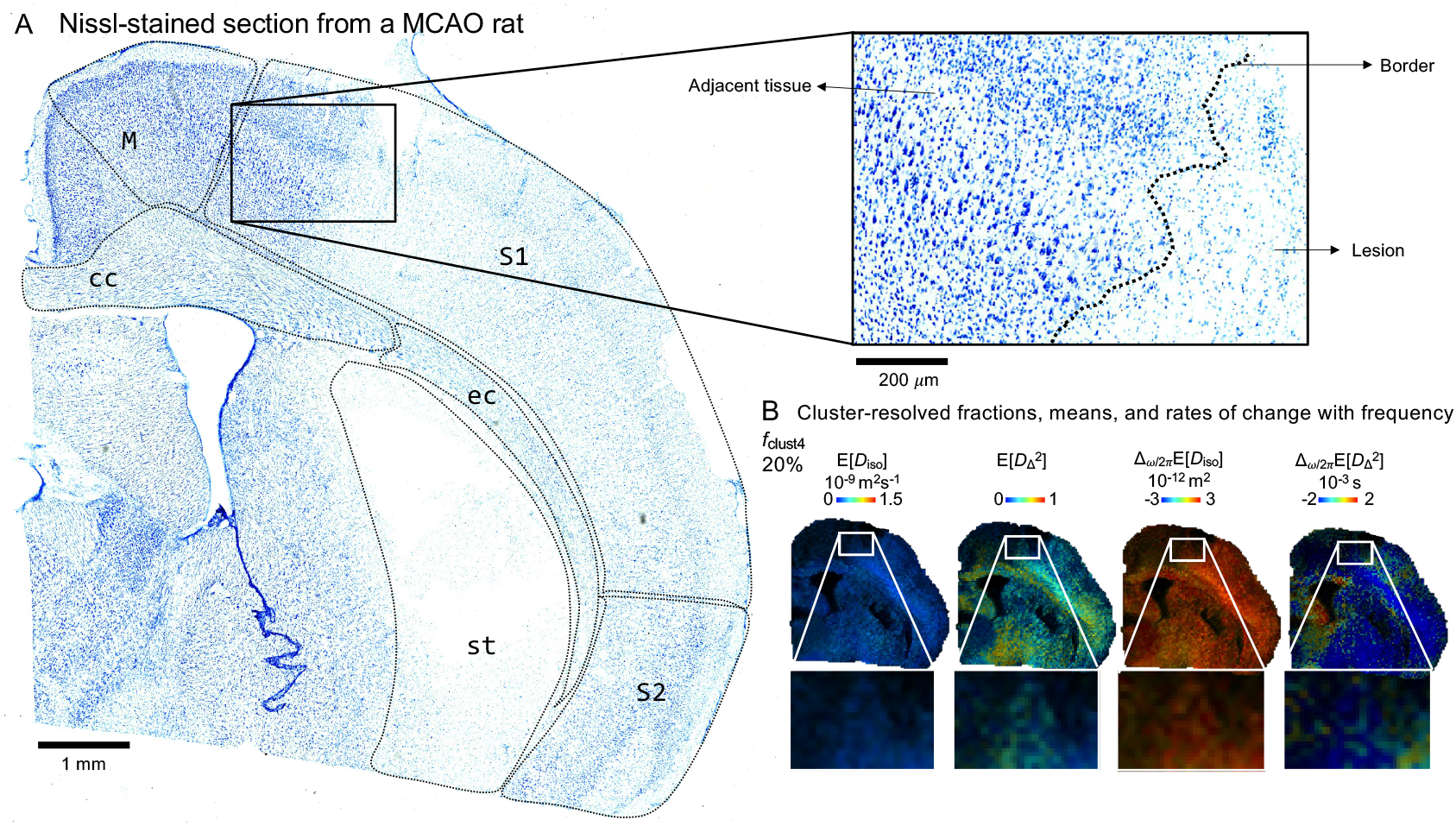
Comparison of Nissl-stained section with cluster-resolved parametric maps from a reprasentative MCAO rat. **A)** A Nissl-stained section of a representative MCAO brain showing the overall tissue damage, with the black arrow indicatin ischemic lesion. **(B)** Cluster-resolved ωDTD maps of f_clust4_ of the same brain, which represent the fraction associated with the lesion he high-magnification of the Nissl photomicrograph **(A)** and close-ups of the MRI parameter maps **(B)** display the same lesio boundary region in the somatosensory cortex. The closer view of the Nissl shows the shrinkage and loss of neurons in the lesion typica in ischemia associated neurodegeneration. The labels are M = motor cortex, S1 = primary somatosensory cortex, S2 = Secondar somatosensory cortex, st = striatum, cc = corpus callosum, and ec = external capsule.

The results of the RF regression fitted for the histological parameters in their original scale are summarized in **Table 1**. The R values for the CV LOO were positive for all the histological parameters and MRI parameter set combinations, indicating that the models captured a relationship between MRI and histology. **Table 1** also shows an increase in the performance metrics for all histological parameters associated with the change of the MRI parameter set from conventional DTI to *ω*DTD. The variation of these metrics was minimal inside the *ω*DTD parameter sets. Based on the *R*2 and *Q*2 values, the model fits (training error) were more accurate than the out-of-sample predictions suggesting overfitting. The best model of cluster-resolved *ω*DTD for number of cells explained 73% of the variance in the data. The improved performance of the model associated with the changing of MRI parameter sets from conventional to *ω*DTD can be seen in supplementary figure S10.

**Table 1:** Performance metrics for the non-linear RF regression analysis. The mean and variance of R, Q^2^, MAE and MSE metrics for the LOO CV calculations as well as R^2^ for the final model. The histological parameters are expressed in their original scale. The best predictions for all the histological parameters were achieved with ωDTD cluster-resolved parameter maps as predictors.

| Histological parameters<br>MRI parameter sets | LOO CV |  |  |  |  |  |  |  | Model |
| --- | --- | --- | --- | --- | --- | --- | --- | --- | --- |
| Number of cells<br>Area of nucleus<br>Circularity of nucleus | E[R] | V[R] | E[Q<sup>2</sup>] | V[Q<sup>2</sup>] | E[MSE] | V[MSE] | E[MAE] | V[MAE] | R<sup>2</sup> |
| Conventional<br>DTI | 0.2275 | 0.0156 | -0.1292 | 0.0416 | 103.5989 | 368.4062 | 7.8298 | 0.8398 | **0.4853** |
|  | 0.1289 | 0.0014 | -0.0894 | 0.0066 | 55.4998 | 298.5853 | 5.0937 | 0.4923 | **0.3978** |
|  | 0.1709 | 0.0048 | -0.0244 | 0.0034 | 0.0249 | 1.7777e-4 | 0.0772 | 4.2602e-4 | **0.3510** |
| $\omega$ DTD<br>Per-voxel | 0.2998 | 0.0135 | -0.0429 | 0.0349 | 95.1086 | 194.6423 | 7.4462 | 0.5337 | **0.6624** |
|  | 0.2324 | 0.0051 | -0.0548 | 0.0117 | 53.1447 | 194.5658 | 5.0178 | 0.3939 | **0.6013** |
|  | 0.2902 | 0.0084 | -0.0310 | 0.0533 | 0.0240 | 1.2312e-4 | 0.0803 | 4.8965e-4 | **0.5846** |
| $\omega$ DTD<br>Bin-resolved | 0.3285 | 0.0152 | 0.0074 | 0.0311 | 90.7678 | 226.2984 | 7.2848 | 0.6007 | **0.6998** |
|  | 0.2601 | 0.0051 | -0.0129 | 0.0111 | 51.3140 | 220.1922 | 4.8888 | 0.4501 | **0.6292** |
|  | 0.3054 | 0.0147 | 0.0014 | 0.0620 | 0.0234 | 1.2957e-4 | 0.0798 | 5.490e-4 | **0.6132** |
| $\omega$ DTD<br>Cluster-resolved | 0.2973 | 0.0151 | -0.0058 | 0.0229 | 92.4096 | 244.0897 | 7.3776 | 0.5053 | **0.7289** |
|  | 0.2168 | 0.0060 | -0.0172 | 0.0043 | 51.9607 | 264.9770 | 4.8845 | 0.5087 | **0.6371** |
|  | 0.2594 | 0.0131 | -0.0397 | 0.0995 | 0.0245 | 1.4979e-4 | 0.0855 | 9.2021e-04 | **0.6086** |

The permutation test results comparing DTI against the three *ω*DTD methods are summarized in Table 2. Across all metrics and histological parameters, *ω*DTD methods showed significant superior predictive performance over conventional DTI in 22 out of 36 comparisons (p<0.05), with particularly strong effects for number of cells prediction where all three *ω*DTD variants significantly outperformed DTI (p<0.05 across *R, Q*2, MSE, and MAE). For area of nucleus, 9 out of 12 comparisons significantly favored *ω*DTD, with per-voxel and bin-resolved methods showing consistent significance across metrics. Circularity of nucleus showed the weakest MRI-histology relationships overall, with only 3 significant comparisons. P-values showing significantly superior DTI were rare, while p-values showing superior *ω*DTD dominated across all metrics. The rest of the permutation testing results can be found in Table S3.

**Table 2:**
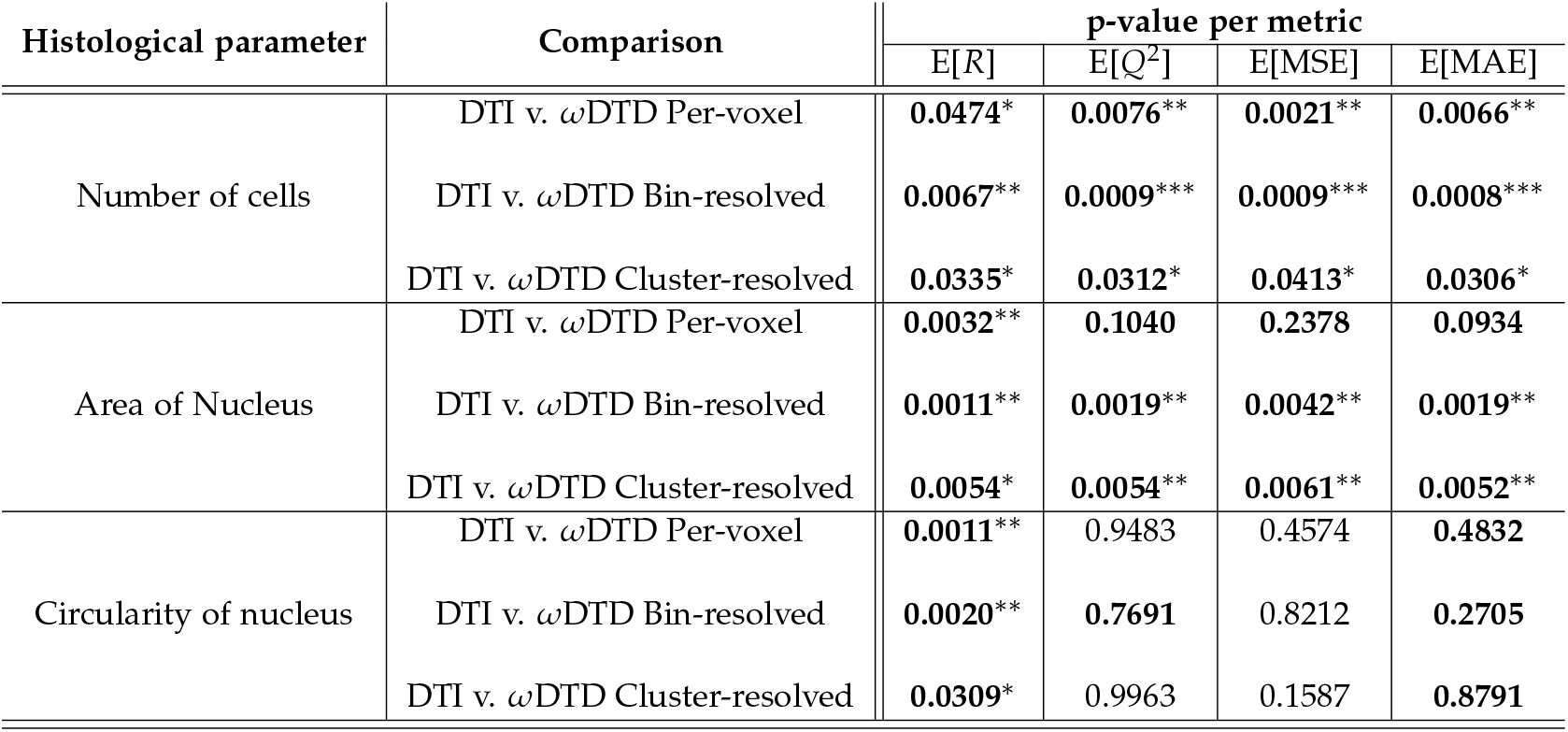
Two-sided permutation test p-values on random forest LOO CV metrics. **Bold** indicates instances where ωDTD methods performed better compared to DTI. ^∗^p<0.05, ^∗∗^p<0.01, ^∗∗∗^p<0.001

The normalised global SHAP mean absolute values for the top ten features of the *ω*DTD cluster-resolved mean and rates of change per frequency maps can be seen in figure 6. The E[*D*_iso_] of *f*_clust1_ emerged as the dominant feature, followed by E[*D*_iso_] of *f*_clust4_ and 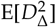 of *f*_clust1_. The different clusters of E[*D*_iso_] occupied six of the top ten positions in global feature importance. And while E*D*_iso_ of *f*_clust1_ ranked highest overall, E*D*_iso_ of *f*_clust3_ and *f*_clust4_ demonstrated the greatest consistency within the top ten global features.

**Figure 6:**
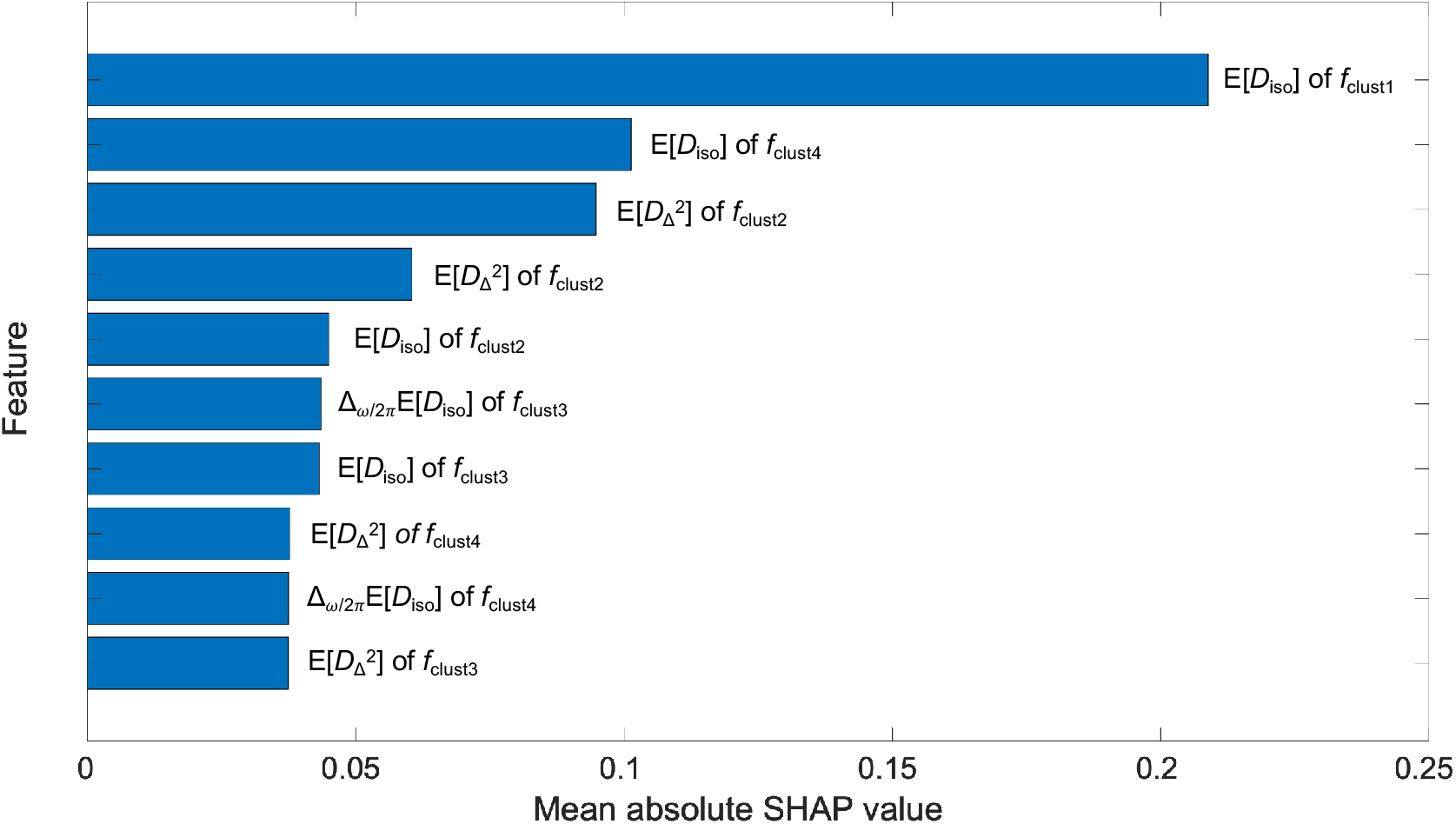
Normalised global SHAP mean absolute values for random forest feature importance. Global SHAP values were calculated in the CV LOO folds for out-of-loop subjects for the ωDTD cluster-resolved mean and rates of change per frequency maps.

The predicted histological parameter maps based on the *ω*DTD cluster-resolved MRI parameter set is shown in **Figure 7**. There was no significant under-estimation of values when comparing the original histological parameter maps (**Figure 7A**) and to predicted ones (**Figure 7B**) as shown by the MAE values in **Table 1**. The RF based predictions also retained information on the different anatomical structures, separating between WM and GM as well as the ventricles, which are distinguishable, though they are not valued as null as in the original maps. The model predictions also demonstrated the capacity to separate between the lesion and adjacent tissue. Each histological parameter map of the MCAO brain presented in **Figure 7B** shows subtle differences in lesion boundaries, differences that are replicated in the predicted histological parameter maps (**Figure 7D**). The average difference (in percentages) between the prediction and original show smaller differences in the circularity of nucleus parameter, when compared to the area of nucleus and cellularity parameters.

**Figure 7:**
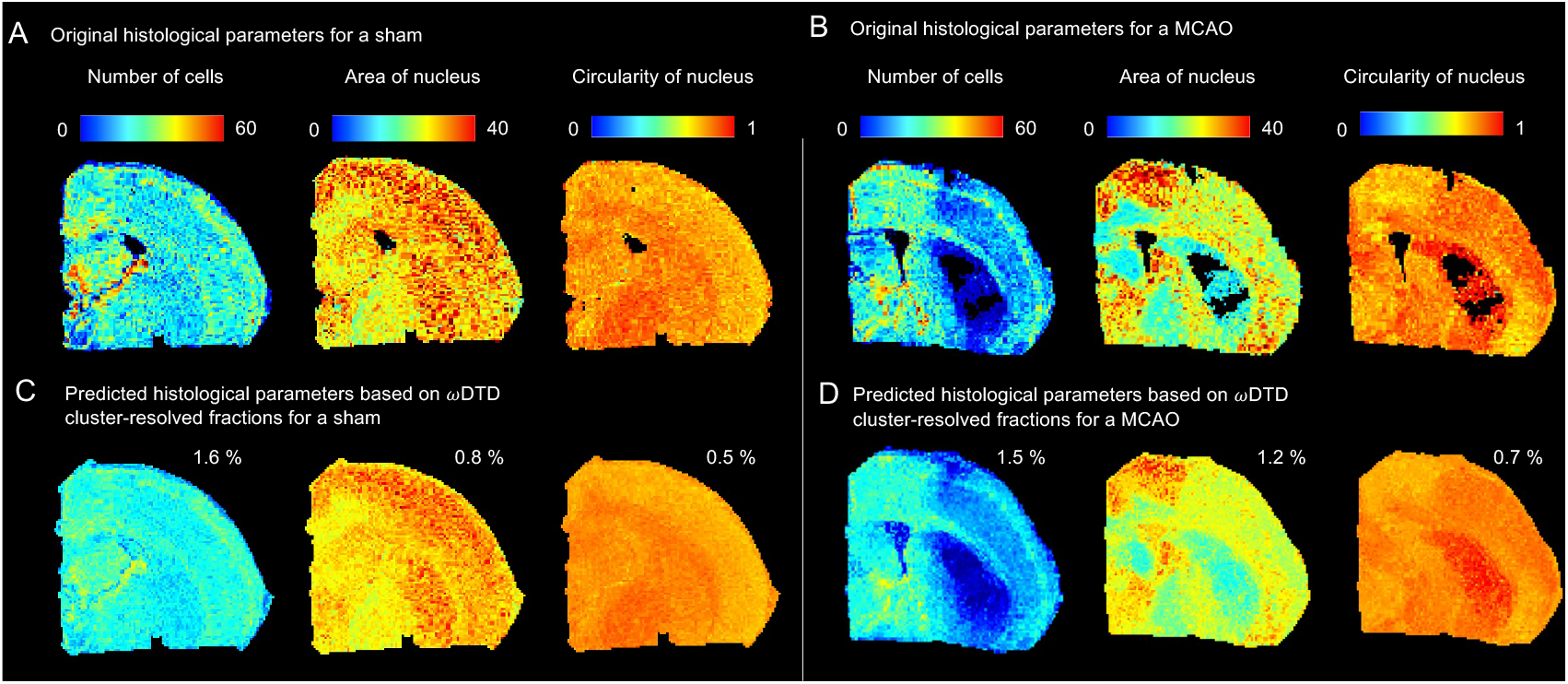
Predicted histological parameters of representative sham and MCAO brains based on non-linear RF regression analysis. Comparison of original downsampled and registered histological parameters (A) and (B) with their predicted counterparts (C) and (D). The predicted histological parameter maps were derived from training a random RF using ωDTD cluster-resolved mean and rates of change with frequency maps. The percentage indicates average difference between the prediction and original.

Overall, the number of cells histological parameter in the MCAO brains show there were less cells in the lesion in comparison to the lesion adjacent tissue. The area and circularity of nucleus parameter maps of the MCAO brains also show decreased nucleus area and increased circularity in the lesion. The distinct boundaries of the lesion within the histological parameters were accurately captured by the predictions.

The distinct details are also reproduced consistently, such as the layer of increased number of cells in the cortex (**Figure 7B** and **D**). The baseline values of nucleus circularity in the secondary somatosensory cortex, an area that was otherwise characterized by a decreased cell number and nucleus area by the other two histological parameters. The original and predictions of all three histological parameters and all the brains are shown in supplementary figure S11.

## DISCUSSION

In this study, we investigated the relationship between *ex vivo ω*DTD parameters and histological parameters in the brain after ischemic damage. We examined the capacity of these MRI parameters in predicting tissue changes seen in the histology. We had four sets of MRI parameters: conventional DTI, *ω*DTD, bin-resolved *ω*DTD, and cluster-resolved *ω*DTD parameters. We found that the *ω*DTD parameters were highly predictive of changes in the cellularity and morphology, including alterations in nucleus size and shape, of cells, more than conventional MRI parameters. Furthermore, we found that some of the histology metrics could be predicted by using a multivariate non-linear regression modeling approach. Standard voxelwise and ROI-based analyses yielded no significant findings, likely due to the small sample size and heterogeneity of the ischemic lesion. Altogether, these results represent a significant step forward in the interpretation of *ω*DTD parameters, showing how they potentially reflect the underlying changes in tissue microstructure after ischemic stroke.

There are numerous studies on using DTI for analyzing ischemic stroke related pathology in both humans and rodents at acute timepoints [52, 53, 54, 55]. Traditionally, DTI is used for detecting the lesion in clinics. MD and FA typically show lower values at an acute timepoint. Our study demonstrates convergent validity in the decrease detected in the lesion in both MD and FA. However, these conventional parameter maps lack specific information on the underlying pathological processes [6, 56]. In studies comparing DTI with more advanced methods, DTI has consistently been found to be less sensitive in detecting subtle microstructural changes and separating specific processes, particularly in cases of early ischemic injury [57, 58, 59].

There are no studies focused on investigating the relationship between *ω*DTD and quantitative histological parameters in the case of ischemic stroke. Our previous studies show that *ω*DTD parameter maps reflect distinct cellular properties [30, 37], which can potentially provide specific insights into the cytopathology through differential lesion segmentation, e.g. the E[*D*_iso_] map in gray matter-like tissues has been linked to cell size. The decrease in values of this parameter seen in the ischemic lesion may be related to the presence of glial cells and neuronal degeneration. The increased values of Δ_*ω*2*π*_E[*D*_iso_] have been linked to increased cellularity in tumors [60, 61]. However, in the case of stroke lesions, which also show increased values in both in bin- and cluster-resolved *f*_clust4_ and *f*_clust6_ maps, this may instead be linked to localized debris such as cellular debris and protein aggregation, creating temporary diffusion barriers. Within the framework of multidimensional MRI (MD-MRI), which encompasses advanced techniques like *ω*DTD, previous studies have shown the stability of both MD-MRI parameters and MD-MRI-derived clusters across subjects [62, 17].

Nissl staining shows a decreased number of cells in the lesion, so another possible reason for the increased values could be changes in the increased extracellular space. The oscillating gradient spin-echo (OGSE) can provide comparison for *ω*DTD [63], although OGSE probes stick-like tensor shapes at different frequencies, while with *ω*DTD we can access a wide range of shapes in the tensor distribution space at different frequencies [30, 35]. We found studies on OGSE in evaluating ischemic stroke. These studies state that necrotic tissue often exhibits increased viscosity or heterogeneous environments due to cellular debris/fragmented membranes, which might amplify restriction effects detectable through OGSE [5, 64, 65]. The decreased values in *f*_clust2_ across all the cluster-resolved *ω*DTD parameters might be reflective also of the breakdown of structural integrity in the lesion, as this cluster-fraction might potentially describe the dense intracellular networks of neurons [12, 17]. The increased values in 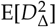 of *f*_clust4_ could further support an increase in structural disorder in the lesion indicating the potential of heterogeneous environments in the lesion.

The coefficient of determination *R*2 is used to evaluate the model’s performance on training data. Based on the *R*2 values, our models can explain between 35% - 70% of the variance present in the data depending on the histological and MRI parameter set. We also found that the number of cells parameter was more substantially associated with the MRI data than the area and circularity of the soma. And even though, on average, the difference between the original and predicted circularity of nucleus parameter measured in percentages was the smallest, the small value range of 0 to 1 means even small errors impact the variance and affect the *R*2. The consistently positive correlation *R* in the CV LOO shows that our models repeatedly capture a relationship between the input MRI parameters and the target histological parameters. Additionally, the jump of increasing values of *R* from conventional to *ω*DTD parameters indicate that the latter MRI parameters are more representative of the histological target parameters. *Q*2 is commonly used in context of CV to assess the model’s ability to generalize unseen data. These *Q*2 values were understandable as considering the highly complex nature of biological data, as the distributions in the target variable values in the new unseen data can be very different compared to the training data. This is especially true in our dataset, as the MCAO lesion is very heterogeneous in size, location and severity. RFs are not especially prone to overfitting as they include regularization through bagging [66], but the grouped nature of the data (i.e. several datapoints from the same animal) might not be optimal for bagging based regularization as it somewhat violates the independence assumption of data points.

We also calculated the MAE and MSE values for the LOO CV. The spread of both MSE and MAE also suggests some variance in model performance across the folds of the CV. Especially, the variance of MSE indicates that there are large errors in certain CV folds, which are penalized heavily due to the squaring of errors. This implies that the model performs inconsistently across the test folds, which suggests differences between individual animals. The difference in the MSE values associated with the MRI parameter sets shows improved and more consistent performance across the CV folds with less large errors when moving from conventional to *ω*DTD parameter sets. This phenomenon along with the increase of *R*2 values is reasonable, as the performance of a RF model is tied to the information content, but also the amount, of data fed into the algorithm. The increased amount of data helps reduce variance while providing the RF more opportunities to identify robust and generalizable patterns. More descriptive data, i.e. data more strongly related to the target variable, on the other hand allow the decision trees to make more confident splits leading to improved predictive power of the ensemble.

The permutation test results comparing conventional DTI against the three *ω*DTD methods provide robust statistical evidence that multidimensional diffusion MRI parameters yield systematically higher predictive performance across LOO CV folds. For number of cells, all *ω*DTD variants significantly out-performed DTI across all four metrics, confirming the markedly stronger relationship between MRI features and neuronal density quantification. Area of nucleus showed a similar pattern with 9 out of 12 significant comparisons favoring *ω*DTD, whereas circularity exhibited the weakest and most inconsistent MRI-histology coupling with few significant differences. The predominance of comparisons indicating superior *ω*DTD for *R* and Q2 suggests that *ω*DTD’s microstructural sensitivity enhances correlation-based generalization while giving superior absolute prediction accuracy. This inter-subject permutation testing framework validates that the observed performance gains from conventional to *ω*DTD parameter sets reflect true methodological improvements rather than random variation.

The dominance of E[*D*_*iso*_] clusters in the global SHAP rankings underscores the model’s reliance on isotropic diffusivity as a primary biomarker of stroke-related tissue damage across the whole brain. The leading role of E[*D*_*iso*_] of *f*_clust1_ could be explained by this cluster’s substantial representation (∼20% of voxels per animal) along with its excellent lesion contrast. This suggests the model captures both cytotoxic edema in the lesion core and possibly the decrease of organization. E[*D*_*iso*_] of *f*_clust3_ and *f*_clust4_ provide complementary information on the overall data: *f*_clust3_’s oblate, frequency-sensitive profile might reflect disrupted neuronal processes, while *f*_clust4_ delineates the primary pathological compartment i.e. edema. Across the entire dataset, E[*D*_*iso*_] along with these three clusters clearly emerge as the primary drivers of model predictions.

There are some limitations in this study that should be considered. First, for future studies, the sample size should be increased to extract more reliable statistical information. Secondly, the MCAO model produces a quite variable ischemic lesion, both in size and location. This heterogeneity also brought challenges to conventional statistical analyses. Third, the acute 24-h timepoint is a quite short period of time for any secondary changes in distal tissue to be detectable. This timepoint also provides a very limited window for the MRI to show the tissue response to the neuroglobin treatment, particularly given that the treatment’s goal at the 24-hour timepoint is primarily neuroprotection [22]. Additionally, infarct maturation beyond 24 h (e.g., toward 72 h or 7 days) could alter *ω*DTD, reducing insight into temporal robustness.

As a fourth, we have only compared conventional DTI to the *ω*DTD parameters, without including other advanced post-processing methods, such as soma and neurite density imaging (SANDI) or the standard model [67, 68, 69]. Single diffusion encoding (SDE) data acquired from the same tissue samples as advanced diffusion measurements would enable more rigorous comparisons. This represents an important direction for future work and will be addressed in subsequent studies. Fifth, histology-to-MRI registration was performed using rigid similarity transforms, which may not fully account for local tissue deformations caused by ischemia, fixation, and sectioning. More advanced non-linear or blockface-guided registration strategies with quantitative target registration error assessment could improve the accuracy of voxel-wise MRI-histology correspondence. Sixth, Nissl staining was the only histology method included in our study, and other stains could add more information about the complex changes occurring in the tissue after ischemia, e.g. myelin damage or inflammation. Finally, we used the automated cell detection with default settings provided by QuPath to extract quantitative data from the histology. These algorithms require further development to increase their accuracy and extract more valuable parameters of tissue damage.

Future steps will be to expand the analysis to fully 3D, for both the MRI and histology. Our frequency-dependent MRI data was 2D due to technical constraints of our sequence at the time, which has since been formatted for 3D acquisition for *ex vivo* and *in vivo*. The latest MD-MRI *in vivo* protocols exist for both preclinical and clinical settings, achieving whole-brain coverage in 20-40 minutes depending on parameter selection, *ω*-ranges, and inclusion of T_1_/T_2_ variation. These example protocols for **D** −*T*_1_ −*T*_2_ and **D**(*ω*) −*T*_1_ −*T*_2_ presented in [17, 70, 37], establish that streamlined MD-MRI approaches, such as *ω*DTD, are achievable despite key constraints of scan time and gradient demands. A feasible 40-minute **D**(*ω*) - *T*_1_ - *T*_2_ acquisition of the human brain employs varying b-values (0–3×109 s/m−2), normalized anisotropy *b*_Δ_ (–0.5, 0, and 1), varying repetition time *τ*_*R*_ (620–7000 ms), and echo time *τ*_*E*_ (40–150 ms), orientation (*θ,ϕ*) and centroid frequency *ω*_cent_/2*π* within 10–90% percentile (4–9 Hz). We are additionally in the process of analysing recently acquired longitudinal *in vivo* preclinical **D**(*ω*) −*T*_1_ −*T*_2_ data to investigate the temporal effects of both the neuroglobin treatment and changes in **D**(*ω*) −*T*_1_ −*T*_2_ parameters and −derived clusters for a future publication.

We will also include other stains to help better interpret the *ω*DTD parameters in terms of histological changes and to understand the relationship between methodologies. With increased histological data, it would be possible to extract more quantitative parameters for our analyses. Additionally, for the analysis, hyperparameter tuning and grid searches for determining the best possible combinations for the RF regression model could also improve the performance [71]. This could improve the robustness of our model and ability to generalize. Incorporating deep learning elements and algorithms could also open new avenues and perspectives in predicting tissue microstructural changes based on imaging outcomes during pathological conditions.

## CONCLUSION

In conclusion, our findings show that *ω*DTD better predicts the changes in quantitative tissue microstructure parameters than conventional DTI after ischemic stroke. And, although our models could not perfectly explain the data, this study indicates that *ω*DTD can open new avenues in the evaluation of ischemic stroke by offering higher sensitivity to the changes associated with ischemic injury, even in its earliest stages. By capturing subtle cellular level alterations in tissue viability, composition, and microstructure, *ω*DTD can enhance our understanding of ischemic stroke progression and improve clinical decision-making along with facilitating better monitoring of treatment responses.

## DATA AND CODE AVAILABILITY

The md-dmri MATLAB toolbox for preprocessing and Monte-Carlo data inversion, along with codes for Gaussian mixture model clustering are freely available at https://github.com/UEF-Multiscale-Imaging. The data used in this study is available upon request.

## ACKNOWGELEDGEMENTS

The computational analyses were performed with the servers of the Bioinformatics Center, University of Eastern Finland, Finland, and the supercomputers at CSC — IT Center for Science, Finland. This study was carried out with the support of the Kuopio Biomedical Imaging Unit, University of Eastern Finland, Kuopio, Finland (part of the Finnish Biomedical Imaging Node, Euro-BioImaging), and the technical and human support provided by the *Centro de Instrumentación Científico-Técnica* (CICT) and by the *Centro de Producción y Experimental Animal* (CPEA) — *Servicios Centrales de Apoyo a la Investigación* (SCAI) — Universidad de *Jaén (UJA, MICINN, Junta de Andalucía, FEDER*).

## SUPPLEMENTARY MATERIAL

**Figure S8:**
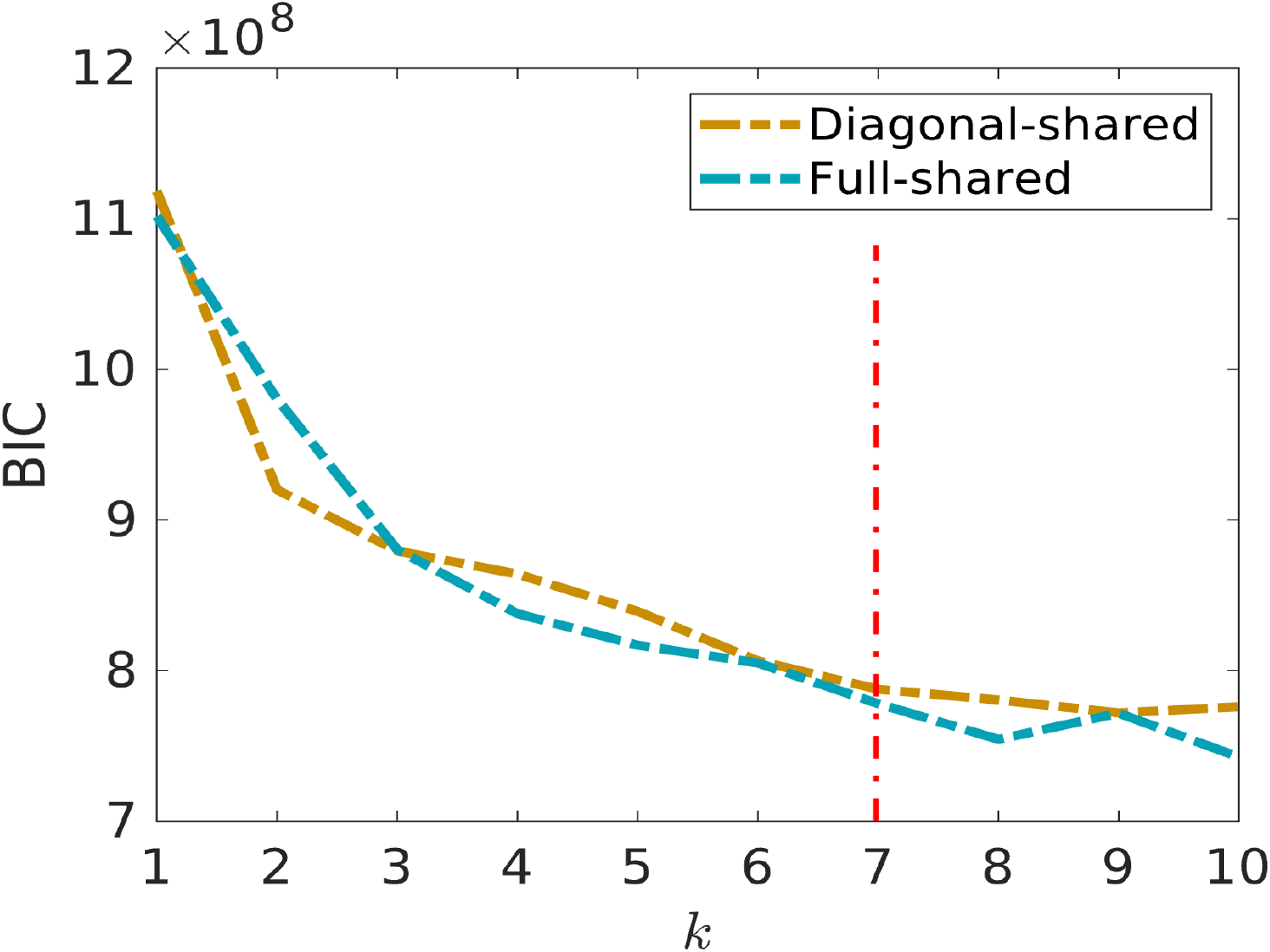
Bayesian information criteria (BIC) in aiding the selection of optimal value for k in unsupervised clustering of D(ω) distributions. The red dotted line indicates the optimum k, chosen as the point of stabilization. Diagonal and full shared refer to the allowed shapes of the clusters.2

**Figure S9:**
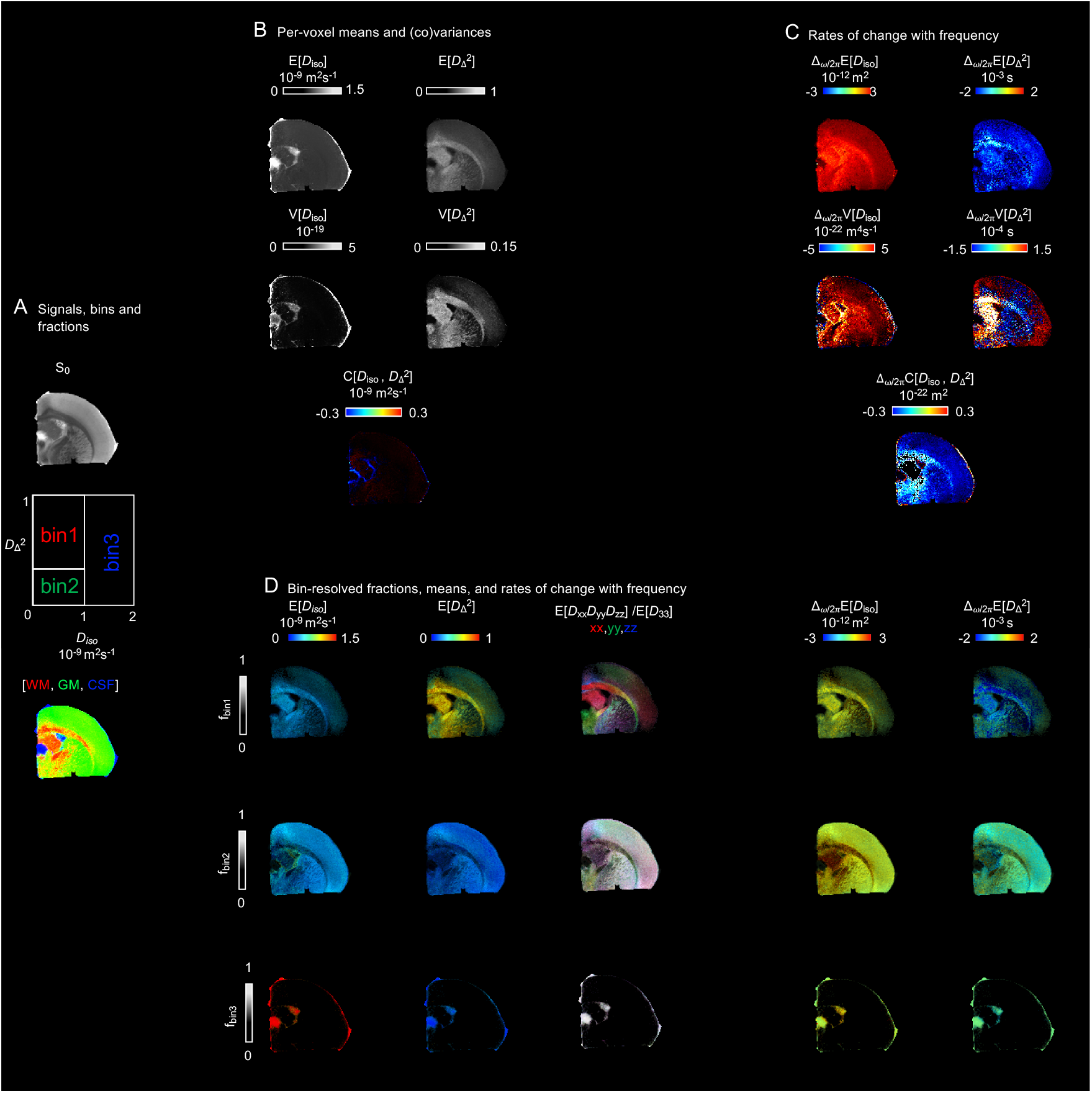
Parameter maps derived from the per-voxel D(ω) distributions for a representative sham brain. **(A)** S0 map in grayscale, the diagram depicting the division of the 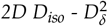 projection into three bins (bin1, bin2, and bin3) and the resulting signal fractions (fbin1, fbin2, and fbin3) coded into RGB color. (B) The per-voxel means E[X], variances V[X], and covariances C[X] at a selected encoding frequency at ω/2π = 34.4 Hz. (C) Parameter maps of the rate of change with frequency Δ_ω/2π_ of the per-voxel means, variances, and covariances of D_iso_ and 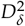 calculated with ω/2π = 34.4-115 Hz. (D) The bin resolved maps of E[X] and Δ_ω/2π_E[X]. The brightness of the bin-resolved maps reflects signal fraction, while the color indicates parameter values.

**Figure S10:**
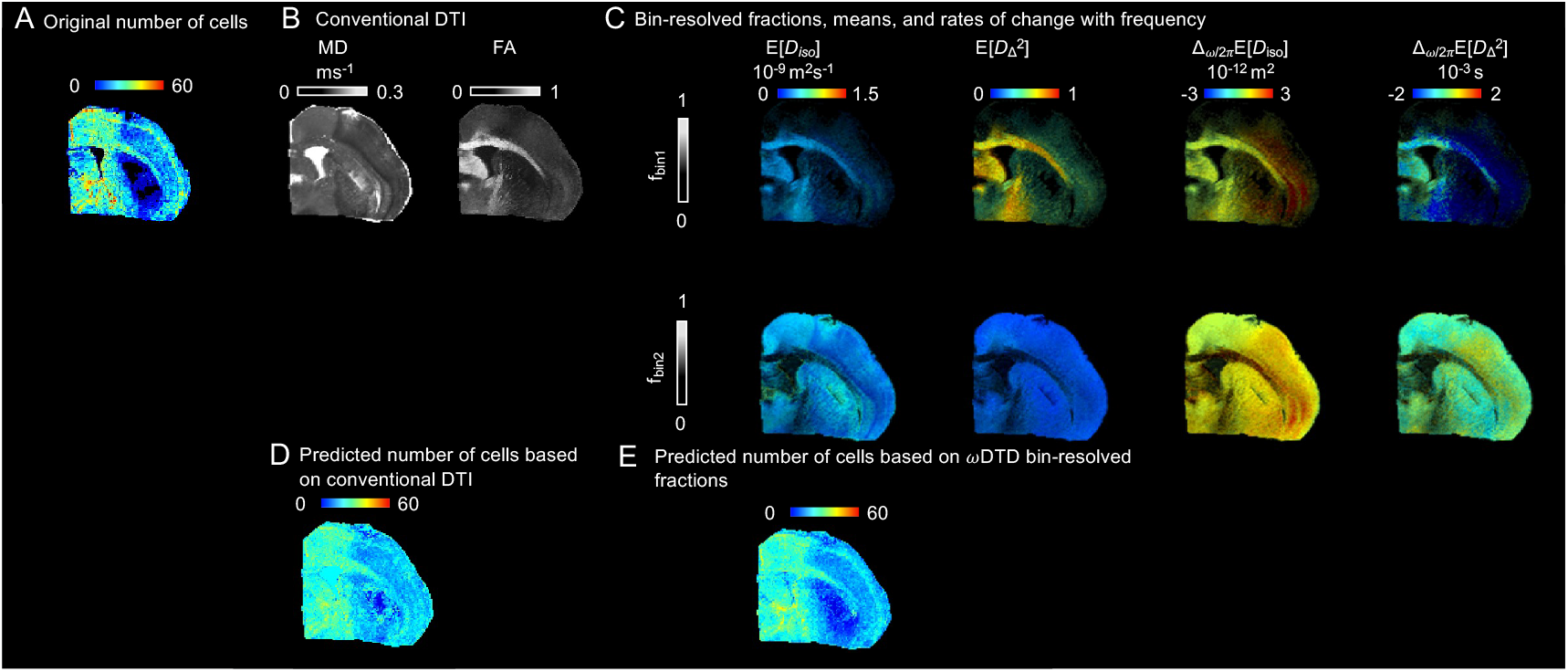
Example of the difference between the non-linear regression predictions for number of cells per voxel histological parameter based on two different MRI parameter sets. The two MRI parameter set are shown in (B): Conventional MRI and (C): ωDTD bin-resolved mean and rates of change per frequency maps (equations (4),(6),(7), and (8)). The predicted number of cells per voxel based on the conventional MRI parameter set (D) demonstrates near identical lesion details as can be seen in the MD map in (B), highlighting a scenario where the regression has clearly failed to capture the nuances of the histological tissue structure. This prediction also lacks the distinct details of the lesion as seen in the original map in (A), in addition to underestimating the values. In comparison, the predicted number of cells per voxel based on the ωDTD bin-resolved fractions (E), resembles more closely the original map in (A) both in lesion details and values, while not directly copying information from any singular MRI parameter.

**Figure S11:**
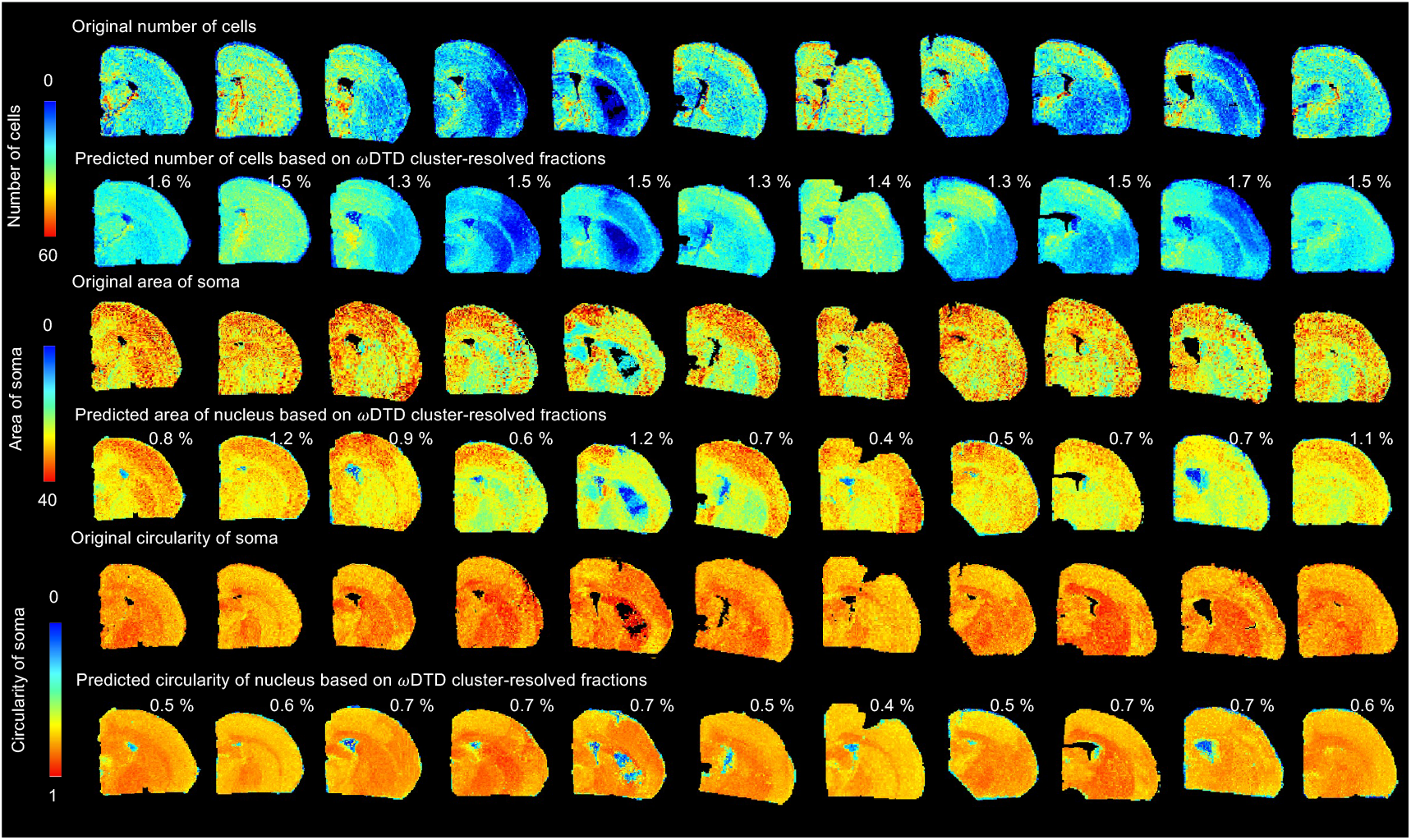
Predicted histological parameters based on non-linear RF regression analysis for all brains. Comparison of original downsampled and registered histological parameters (rows) for the sham (first column) and all MCAO brains (columns 2-11) with their predicted counterparts. The predicted histological parameter maps were derived from training a RF model using Ωdtd cluster-resolved mean and rates of change with frequency maps. The figure displays data across all the brain samples and histological parameters, providing a comprehensive comparison between the observed and predicted histological features. The percentage indicates average difference between the prediction and original.

**Table S3:**
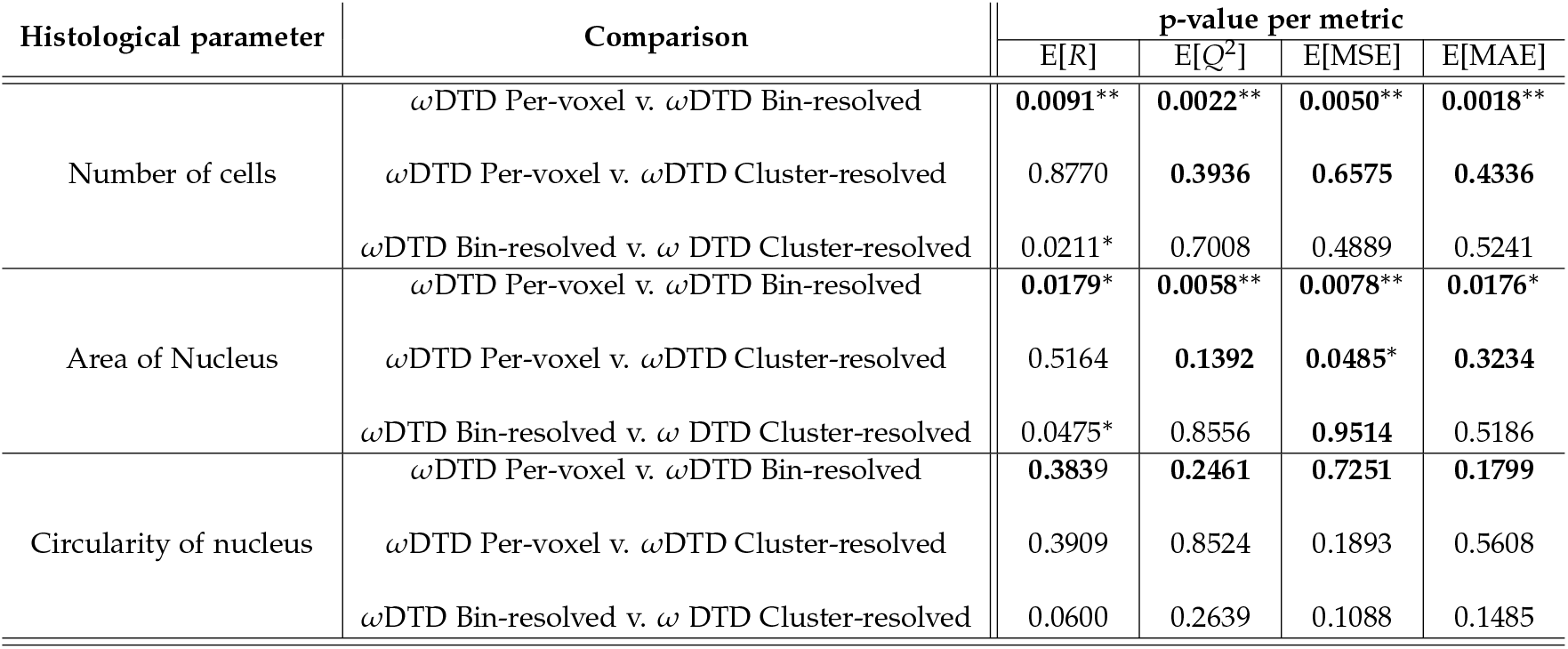
Two-sided permutation test p-values on random forest LOO CV metrics. **Bold** indicates instances where the 2^nd^ ωDTD method performed better compared to the 1^st^ ωDTD method. *p<0.05, **p<0.01, ***p<0.001

